# Reprogramming the neuronal secretory and metabolic machinery using correctors for Alzheimer therapy

**DOI:** 10.1101/2024.11.09.622805

**Authors:** Cho Won Jin, Yutong Shang, Sushmita Patil, Xin Qi, Bhanu P. Jena

## Abstract

Alzheimer disease (AD) is a neurodegenerative disorder characterized by memory loss and personality changes, leading to dementia. The primary cause of the cognitive decline that characterizes AD is the loss of neurons, hypothesized as being due to secretory defects in neurotransmitter release and synaptic plasticity. There is growing evidence that in Alzheimer there is impaired neuronal metabolism due to an increase in free radicals and mitochondrial fission, leading to loss in ATP synthesis required for neurotransmission. Therefore, impaired metabolism in neurons may lead to the observed defects in neurotransmitter release, resulting in the loss of neuronal function and connections in regions of the brain involved in memory and reasoning. In our earlier studies, peptide inhibitor targeting oligomeric ATAD3A, a protein essential for mitochondrial fission and bioenergetics, were found to partially rescue from AD, both in AD neurons and AD mice. In the current study in AD neurons, neuronal porosome reconstitution combined with either a linear or circular peptide inhibitor of ATAD3A oligomers, was able to restore AD neurons to near normal levels of viability and greatly reduce oxidative stress in mitochondria. Additionally, the peptide and a small molecule considered safe by the FDA and used as a food additive, were also able to greatly increase viability in AD neurons and provide reduction to near normal levels of mitochondrial oxidative stress. Collectively, these results demonstrate great promise as secretory and metabolic correctors for AD therapy.

## INTRODUCTION

Alzheimer’s disease (AD) is a neurodegenerative disorder characterized by memory loss leading to dementia. It is estimated that in the United States nearly 6.2 million individuals are impacted by AD dementia, and over 50 million globally. AD is increasing at an alarming pace, projected to double by 2050. According to the *World Alzheimer*’*s Report*, the total estimated annual worldwide health care costs for persons with AD has reached almost a trillion-dollars [1]. Secretory defects in neurotransmitter release and synaptic plasticity, synaptic loss, and mitochondrial dysfunction, have been recognized as early events in AD pathogenesis [2]. Brain regions, especially the entorhinal, temporal and frontoparietal cortex, the hippocampus and the subcortical nuclei are primarily affected [3]. There is growing evidence that in Alzheimer’s there is impaired neuronal metabolism due to an increase in free radicals and mitochondrial fission, leading to loss in ATP synthesis required for neurotransmitter release [4-6].

### Neurotransmitter Release

Porosomes are secretory portals at the cell plasma membrane where secretory vesicles transiently dock and fuse to expel a precise amount of intra- vesicular contents from the cell during secretion [7,8]. In neurons, porosomes are 15nm cup- shaped lipoprotein structures at the presynaptic membrane, composed of nearly 40 proteins [9]. A number of porosome proteins have previously been implicated in neurotransmission and neurological disorders, attesting to the crosstalk between porosome proteins and their coordinated involvement in release of neurotransmitter at the synapse. In Alzheimer’s, levels of porosome proteins CNPase (2,3-cyclic nucleotide phosphodiesterase) and the heat shock protein 70 (HSP70) are found to increase, while the levels of dihydropyrimidinase-related protein-2 (DRP-2) decrease [10]. Decreased levels of CNPase have been observed in the frontal and temporal cortex of patients with AD [11]. Similarly, porosome proteins SNAP-25 and synaptophysin are significantly reduced in neurons of patients with Alzheimer’s disease [12-14]. Mice that are SNAP-25 (+/-) show disabled learning and memory, and exhibit epileptic like seizures [15]. These results support the conclusion that, alteration of one porosome protein impacts others within the complex, resulting in impaired porosome- mediated secretion. This finding is similar to our recent studies on human bronchial epithelial (HBE) cells which showed that the ΔF508 cystic fibrosis transmembrane conductance regulator (CFTR) mutation in HBE cells, affects nearly a dozen porosome proteins including CFTR within the porosome complex. Therefore, the reprogramming of the porosome secretory machinery into the cell plasma membrane of Cystic Fibrosis (CF) cells, was able to rescue from CF [16]. Similarly, we hypothesized that reconstitution of the 15 nm normal neuronal porosome complex in Alzheimer’s neurons would overcome secretory defects in neurotransmitter release.

### Neuronal Energy Metabolism

It is widely accepted that the mitochondrial production of reactive oxygen species (ROS) contributes to the detrimental alterations in the etiology and/or progression of many pathological conditions, including brain neurodegeneration [17]. Studies report that flavonoids can protect cells from different insults that lead to mitochondria- mediated cell death, and epidemiological data further show that some of these compounds attenuate the progression of diseases associated with oxidative stress and mitochondrial dysfunction [18]. Flavonoids are low molecular weight phenolic compounds, displaying significant ROS scavenging capability, including other cellular antioxidant effects [19-21], and are hence ideal for the protection of brain neurons from the buildup of free radicals generated by the mitochondria leading to mitochondrial fission in Alzheimer’s.

ATAD3A is an ATPase family AAA-domain containing protein 3A found at mitochondrial contact sites. It spans the inner and outer mitochondrial membranes, with its N-terminal domain in the cytosol and C-terminal domain in the matrix [22-25]. ATAD3A regulates mitochondrial fission-fusion dynamics. Although global ATAD3A knockout (KO) is embryonic lethal [26], selective ATAD3A KO in mice causes mitochondrial fragmentation and mitochondrial bioenergetics failure, resulting in cell death and tissue damage [27,28]. ATAD3A mutation is also associated with axonal neuropathy and spastic paraplegia [29,30]. Therefore the proper function of ATAD3A is crucial for mitochondrial activity, integrity and cellular survival. In AD, ATAD3A undergoes oligomerization, resulting in Drp1-mediated mitochondrial fragmentation, leading to neurodegeneration [31]. Heterozygous knockdown of ATAD3A in AD transgenic mice abolishes ATAD3A oligomerization, reduces amyloid accumulation and neuroinflammation, and improves long-term memory [31]. These data further support the idea that aberrant ATAD3A oligomerization is a key pathogenic factor inducing AD neuropathology and further validates ATAD3A as a drug target.

### Peptide inhibitor of ATAD3A oligomer

We have developed DA1, a synthetic peptide that binds to ATAD3A and reduces ATAD3A oligomerization, which in turn decreases Drp1/ATAD3A binding that occurrs during AD progression [31,32]. DA1 treatment reduces mitochondrial bioenergetic defects, and improves the survival rate of neurons exposed to toxic Aβ *in vitro*. DA1 penetrates the blood-brain-barrier, and is nontoxic with minimal effects on the immune response [31,31]. These physiological and pharmacodynamic properties make DA1 a promising candidate for AD therapy. We have tested [31] the *in vivo* efficacy of DA1 in 5XFAD AD transgenic mice using a subcutaneous (SQ) Alzet mini-pump, delivering either the control peptide TAT or DA1 starting at 6 weeks of age. 5XFAD mice treated with the control peptide exhibited enhanced ATAD3A oligomerization at 6 months of age, which was abolished by treatment with DA1 [31]. Immunohistochemistry of 6-month-old 5XFAD mice brain revealed a significant increase in the density and area covered by amyloid beta (Aβ). DA1 treatment significantly reduced this amyloid load [31]. Furthermore, DA1 decreased immunoreactivity of GFAP (astrocytes) and Iba1 (microglia), suggesting reduced neuroinflammation. Additionally, DA1 treatment from 2 to 8 months of age improved the long-term memory of 5XFAD mice. DA1 treatment had no effect on the behavior of WT mice, further supporting the nontoxic nature of the peptide [31].

### Secretory and Metabolic Correctors for AD Therapy

In the current study, we have further developed the linear DA1 peptide in to a cyclic DA1 analogue that has improved stability and efficacy. We have tested the combinatorial use of the porosome, the small flavonoid molecule Apigenin, and our linear and cyclic DA1 peptides, in treating AD. Results from our study demonstrate great promise in the combinatorial use of secretory (porosome) and metabolic (flavonoid and DA1 peptide) correctors for AD therapy, restoring AD neurons to near normal levels of viability and greatly reducing oxidative stress in the mitochondria.

## 2. MATERIALS AND METHODS

### 2.1 Mouse HT-22 and Neuro2a wt, Neuro2a APPwt and Neuro2a APPswe cell Cultures

Mouse hippocampal HT-22 cells (MilliporeSigma, SCC129) and mouse neuroblastoma cell line Neuro2a cells (ATCC, CCL-131) were cultured in DMEM supplemented with 10% (v/v) heat-inactivated FBS and 1% (v/v) antibiotics (100 unit/mL penicillin, 100 μg/mL streptomycin). Neuro2a cells stably overexpressing human APP wildtype (APPwt) or Swedish mutant (APPswe, K670N and M671L APP, clone Swe.10) obtained from Dr. Gopal Thinakaran (University of Chicago), were cultured as described above. Cells were plated on poly-D-lysine/laminin (P6407, Sigma-Aldrich)-coated culture plates with or without coverslips at an appropriate cell density. All cells were maintained at 37 °C and 5% CO_2_.

### 2.2 Porosome Isolated from Neuro2a wt (Control) for Western Blot analysis and for Reconstitution into Alzheimer’s (Experimental) Neuro2a APPwt, and Neuro2a APPswe cells

*Neuro2a wt* cells were used to isolate porosomes for reconstitution. RIPA buffer (10 mM Tris-HCl, pH 8.0 • 1mM EDTA • 0.5mM EGTA • 1% Triton X-100 • 0.1% Sodium Deoxycholate • 0.1% SDS • 140mM NaCl • Dilute with dH_2_O) containing 0.1mM PMSF, 1mM ATP, and a cocktail of protease inhibitors, was used to lyse the cells. Protein in all fractions was estimated using BCA Protein assay Kit (ThermoFisher Cat. No. 23227, Rockford, IL 61101, USA). A total of 500 μg of the supernatant proteins were incubated with 2 μg of mouse monoclonal antibody raised against SNAP-25 (Santa Cruz, sc-390814) or Syntaxin1a (MilliporeSigma, SAB4502894) overnight, followed by incubation with 40 μl of 50% protein AG Magnetic agarose beads (Thermo Fisher., Cat. No. 78610) slurry for 1 h. The beads were washed with the binding/wash buffer (PBS with 150 mM NaCl) for 5 min at 4 °C with gentle agitation twice, then washed with deionized water once. The proteins were eluted with 100 uL of 0.1 M glycine, pH 2.2 and neutralized with 25uL of 1M phosphate buffer, pH 7.5. Porosome reconstitution into *Neuro2a APPwt and Neuro2a APPswe cells in culture* was achieved by exposing 0.125 μg/mL porosomes isolated from *Neuro2a wt* cells.

### 2.4 Control TAT and DA1 Peptides

Control peptide TAT and DA1 peptide (Product number P103882, Lot# 0P082714SF-01) were synthesized at Ontores (Hangzhou, China). Their purities were assessed as >90% by mass spectrometry. Lyophilized peptides were dissolved in sterile water and stored at −80 °C until use. DA1 is the first and only inhibitor that selectively binds to ATAD3A, reducing oligomerization, neuropathology, and behavioral deficits in AD mice without affecting wild- type counterparts [31-33]. The peptide is particularly effective under stress or disease conditions, where oligomers accumulate and recruit Drp1. Thus, DA1-like agents offer a promising new therapeutic approach.

### 2.5 Cyclic DA1 peptide analog

In addition to the DA1 linear peptide, we have developed cyclic DA1 analogs with enhanced metabolic stability and improved on-target efficacy for future clinical trials. Following peptide optimization and biochemical screening in AD cell cultures, a cyclic DA1 analog, DW45 has been identified. DW45 display notable stability with a half-life of >289 minutes in a mouse plasma stability assay and >216 minutes in a hepatocyte stability assay, respectively. Moreover, the analogs demonstrates stronger efficacy and on-target engagement in AD neuronal cultures, and was used in the current study.

### 2.6 MTT Assay

MTT (3-(4,5-dimethylthiazol-2-yl)-2,5-diphenyltetrazolium bromide) assay is a colorimetric method that measures cell viability, proliferation, and cytotoxicity. It is based on the reduction of a yellow tetrazolium dye, MTT, into purple formazan crystals by metabolically active cells. Abeta: Ameloid beta; TAT: Transactivator of transcription; DA1 (linear peptide); WX45 (DA1 cyclic peptide)

### 2.7 Western Blot Analysis on cell homogenates and isolated porosomes

Total cell homogenates (TH) and porosomes immuno-isolated (IP) from expanded WT (Control) mouse neuroblastoma cell culture and experimental Alzheimer’s WT-APP and Sew-APP cultures were subjected to SDS-PAGE and Western Blot analysis. 20μg of proteins (TH) and 20μl of isolated neuronal porosomes in Laemmli buffer were resolved in a 12.5% SDS-PAGE, followed by electrotransfer to 0.2-mm nitrocellulose (NC) membrane. The NC membrane was incubated for 1 hour at RT in blocking buffer (5% nonfat milk in PBS [pH 7.4] containing 0.1% Triton X-100 and 0.02% NaN_3_) and immunoblotted for 2 hours at RT with antibodies to SNAP-25 (1:1000, MilliporeSigma, S-9684) and GAPDH (1:3,000, MilliporeSigma, SAB600208). Anti-rabbit horseradish peroxidase secondary antibody conjugates were used (1:5,000, Cell Signaling Technology Cat. No. 7074), and then developed with SuperSignal™ West Femto Maximum Sensitivity Substrate (Thermo Fisher, Cat. No. 34095) and exposed using by ChemiDoc XRT+ image system (Bio-Rad, Richmond, CA94806, USA).

### 2.8 Immunofluorescence Cytochemistry

Cells were grown on coverslips, fixed with 4% paraformaldehyde for 20 min at room temperature, permeabilized with 0.1% Triton X-100 in PBS, and blocked with 2% normal goat serum. The cells were then incubated with the indicated primary antibodies overnight at 4 °C. After washing with PBS, the cells were incubated with Alexa Fluor 488/568 or 405/568 secondary antibody (1:500; Thermo Fisher Scientific) for 2 h at room temperature. The nuclei were counterstained with DAPI (1:10,000; Sigma-Aldrich). Images of the staining were acquired using a Fluoview FV 1000 confocal microscope (Olympus).

## 3. RESULTS AND DISCUSSION

Our earlier *in vivo* studies in AD mice (5xFAD) using the linear DA1 peptide inhibitor of ATAD3A oligomers were shown to partially rescue AD mice from Alzheimer’s [31]. In the current study in AD neurons, both the linear and circular DA1 peptide (WX45) similarly demonstrated a partial rescue from AD both in cell viability and in reduced mitochondrial superoxidase activity (mitoSOX) in Alzheimer’s neurons (Aβ+TAT) [Figure 1, 2]. The WX45 circular DA1 peptide demonstrated to be more effective than the linear DA1 peptide. However as hypothesized, the neuronal porosome reconstitution combined with the DA1 peptide especially the circular WX45 peptide inhibitor of ATAD3A oligomers, was able to restore AD neurons to near normal levels of viability and greatly reduced oxidative stress in mitochondria, demonstrating the potential for therapeutic applications. Additionally, the peptide and Apigenin, a small molecule flavonoid considered safe by the food and drug administration (FDA) and used as a food additive, were also able to greatly increase viability in AD neurons and reduce to near normal levels mitoSOX [Figure 1, 2], also demonstrating its use in AD therapy. Collectively, these results demonstrate great promise in the combinatorial use of the porosome reconstitution secretory reprogramming and the DA1 peptide and Apigenin metabolic correctors, as highly safe and effective AD therapies. In agreement, our immunoblot analysis and immunocytochemistry study of the mouse neuroblastoma Neuro2a control cells, and our APPwt and sweAPP Alzheimer’s cells [Figure 3], demonstrate significant decrease in SNAP-25 immunoreactivity and altered presence of other porosome proteins in both APPwt and sweAPP neurons, confirming earlier reported studies of a decrease in SNAP-25 in patients with Alzheimer’s [13-15]. Consequently, in Alzheimer’s patients there is an increase in levels of SNAP-25 in the cerebrospinal fluid, which has been associated with cognitive decline [34,35]. Upon porosome reconstitution in APP neurons, our immunocytochemistry demonstrates the restoration of SNAP-25 and other porosome-associated proteins [Figure 3] that result in increased cell viability to near normal levels and a decrease in mitoSOX activity [Figure 1].

**Figure 1.**
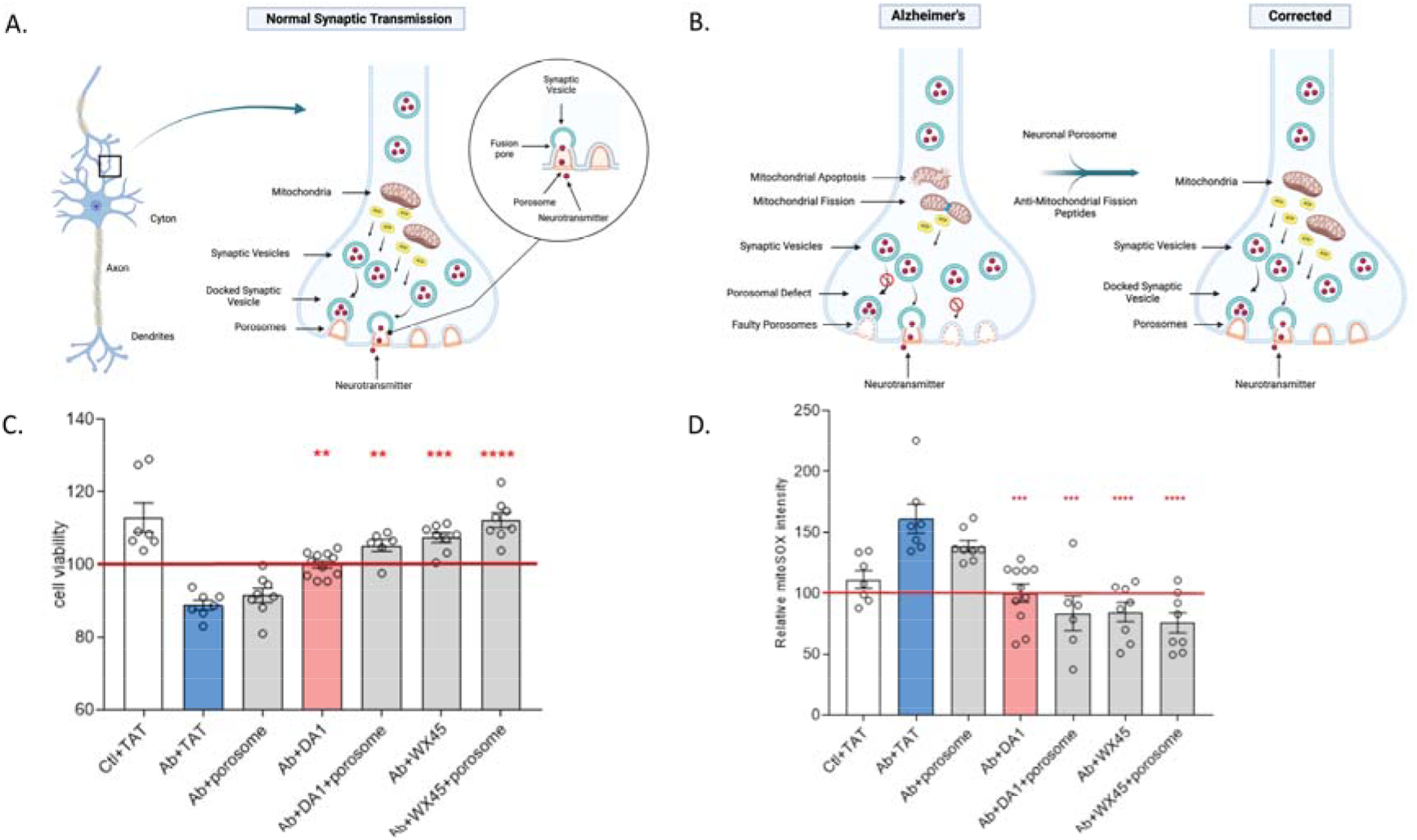
The combinatorial therapy of porosome reconstitution the secretory corrector, and the DA1 peptide inhibitor of mitochondrial fission, the metabolic corrector, restores cell viability and reduces oxidative stress in mitochondria to normal levels in AD neurons. **(A)** Schematic illustration of a synapse at the nerve ending in healthy neurons, demonstrating normal mitochondrial function, energy (ATP) generation, and porosome-mediated neurosecretion. **(B)** Schematic illustration of a synapse at the nerve ending in AD neurons, demonstrating abnormal mitochondrial function resulting in mitochondrial fission, loss in ATP generation, and altered porosome-mediated neurosecretion. All of these defects in AD are seen to be overcome using porosome reconstitution and or the DA1 peptide inhibitors of mitochondrial fission therapy. **(C)** Cell viability is significantly enhanced in Alzheimer’s neurons (Aβ+TAT) by porosome reconstitution either in the presence of the linear or the circular DA1 peptide WX45. HT-22 cells were pretreated with porosome (175ng per well in a 96-well plate) overnight, followed by treatment with oligomeric Aβ_1–42_ peptides (5μM) together with DA1 (linear peptide) or WX45 (DA1 cyclic peptide) (1μM) for 40h. Note that the cell viability in Alzheimer’s neurons following porosome reconstitution and the presence of the circular peptide WX45, demonstrate recovery to healthy neuron levels. Cell viability was measured by MTT assay after 16h of serum starvation. Data represent the mean ± SEM from at least six independent experiments. Statistical significance was determined by one- way ANOVA with Tukey’s post hoc test. **p<0.01; ***p<0.001. ****p<0.0001. **(D)** Porosome reconstitution significantly reduces mitochondrial superoxidase activity (mitoSOX) in Alzheimer’s neurons (Aβ+TAT) in presence of either the linear DA1 or circular DA1 (WX45) peptide. HT-22 cells were pretreated with porosome (175 ng/well in a 96-well plate) overnight. Cells were then pre-incubated with DA1 or WX45 DA1 cyclic peptide (1 μM) for 1 hour, followed by treatment with oligomeric Aβ_1–42_ peptides (10 μM) for 12 hours. Mitochondrial superoxide production was assessed 1 hour after the second peptide treatment using MitoSOX staining. MitoSOX activity in Alzheimer’s neurons following porosome reconstitution demonstrate recovery to much reduced levels than found in healthy neurons in the presence of the circular peptide WX45. Data are presented as mean ± SEM from at least six independent experiments. Statistical significance was determined by one-way ANOVA with Tukey’s post hoc test. ***p<0.001. ****p<0.0001.

**Figure 2.**
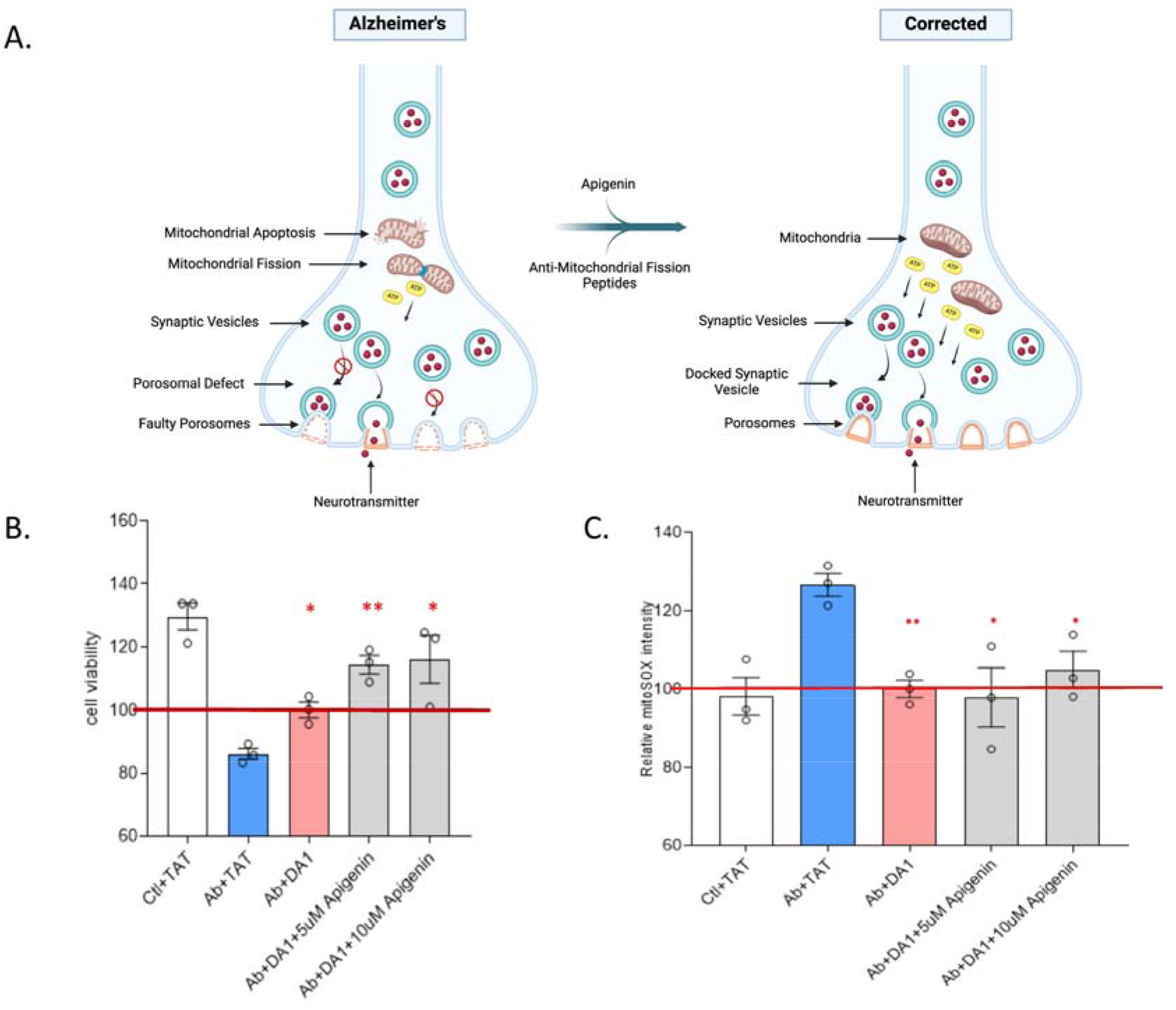
Combination therapy of the small flavonoid molecule Apigenin and the mitochondrial fission inhibiting linear DA1 peptide results in increase cell viability and reduction of mitochondrial oxidative stress to normal levels in AD neurons. **(A Left)** Schematic illustration of a synapse at the nerve ending in Alzheimer’s neurons, demonstrating abnormal mitochondrial function, energy (ATP) generation, and porosome-mediated neurosecretion. **(A Right)** Schematic illustration of a synapse at the nerve ending in AD neurons following a combination of peptide and Apigenin therapy demonstrating normalization of mitochondrial function and corrected ATP generation and neurosecretion. All of these defects in AD are shown to be overcome using Apigenin and the DA1 peptide inhibitor of mitochondrial fission. **(B)** Viability of mouse hippocampal HT-22 neuronal cells exposed to 5μM toxic oligomeric Aβ_1–42_ peptides mimicking Alzheimer’s (Aβ+TAT), is significantly enhanced when treated with 1 μM DA1 linear peptide and 5μM or 10μM Apigenin. Cells were treated with DA1 1μM + Aβ 5μM+ Apigenin (5uM or 10μM). After 24h, cells were washed using serum-free media, and again treated with peptides 1μM + Aβ 5μM + Apigenin (5μM or 10μM) in serum-free medium. Cell viability was measured by MTT assay at 40h (16h serum starved). Data represent the mean ± SEM from at least six independent experiments. Statistical significance was determined by one-way ANOVA with Tukey’s post hoc test. *p<0.05; **p<0.01. (**C)** Mitochondrial superoxidase activity (mitoSOX) in Alzheimer’s neurons (Aβ+TAT) is significantly reduced in the presence of the linear DA1 peptide. However, not much change is observed in the presence of either 5μM or 10μM of Apigenin. Mouse hippocampal HT-22 neuronal cells exposed to DA1 1μM + Apigenin (5μM or 10μM) for 1h and 10μM Aβ (oligomeric Aβ_1–42_ peptides) for 12h. Cells were again treated with DA1 1uM + Apigenin (5μM or 10μM) again 1h before staining for mitoSOX (measure mitochondrial ROS). Data represent the mean ± SEM from at least six independent experiments. Statistical significance was determined by one-way ANOVA with Tukey’s post hoc test. *p<0.05; **p<0.01.

**Figure 3.**
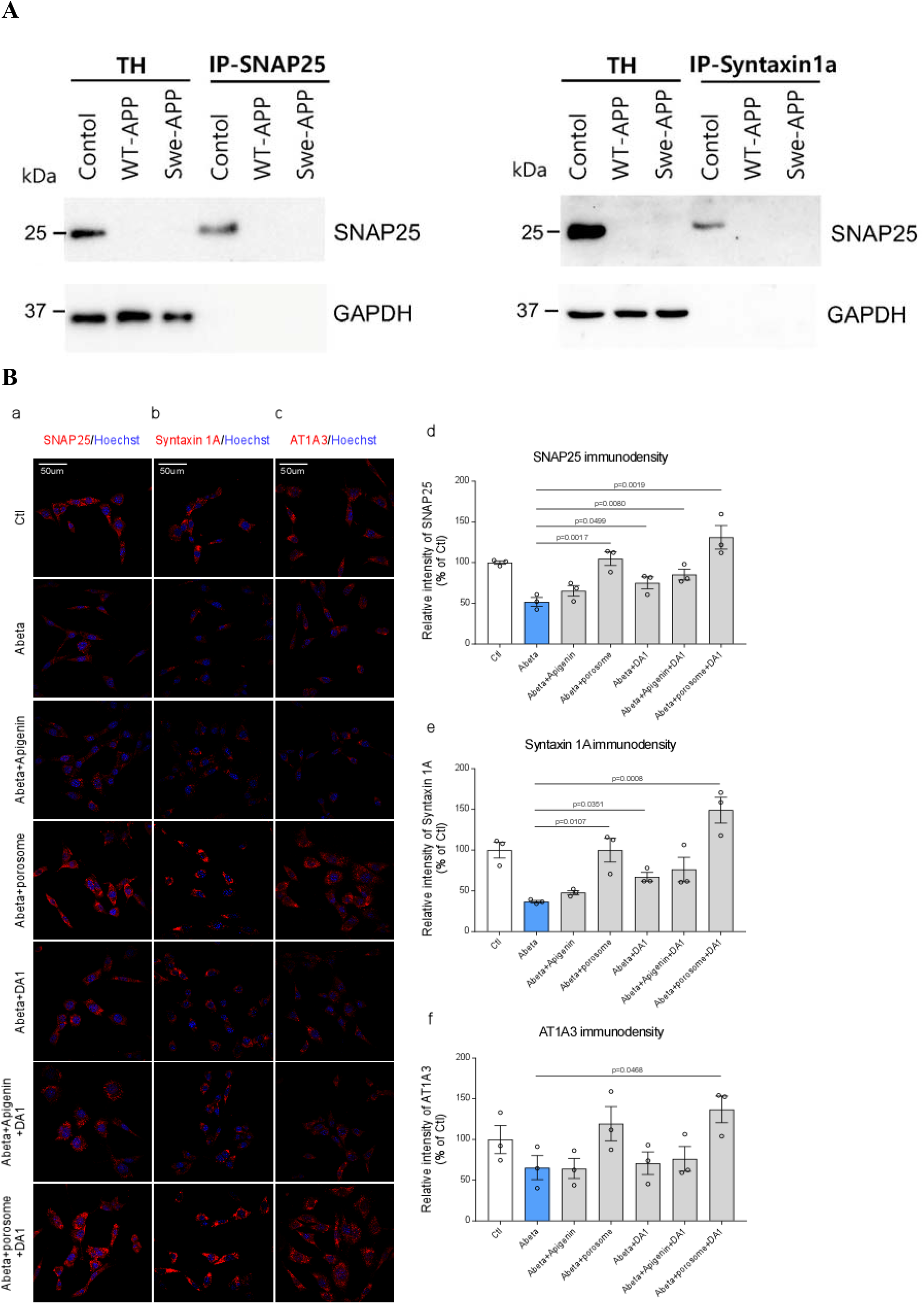
Immunoblot and immunocytochemistry demonstrate depletion of porosome proteins especially the t-SNARE proteins SNAP-25 and Syntaxin-1A, which are restored following peptide and porosome-reconstitution therapy. **(A)** Note that in Alzheimer’s neurons (WT-APP and Swe-APP), there are undetectable levels of SNAP-25 both in total homogenates and in the immunoisolated porosome complex, immunoisolated using the t- SNARE SNAP-25 or Syntaxin-1A antibody. **(B)** Immunocytochemistry demonstrating depletion of porosome proteins SNAP-25, Syntaxin-1A, and Na^+^/K^+^ Transporting ATPase alpha 3 (AT1A3) in Alzheimer’s neurons WT-APP (Aβ), which is fully restored to normal levels following porosome-reconstitution therapy, and partially by the DA1 peptide. The combined exposure to both the porosome and DA1 peptide, further enhances the expression of all three porosome proteins SNAP-25, Syntaxin-1A, and AT1A3. Modest increase in SNAP-25, Syntaxin-1A, or AT1A3 immunoreactivity is observed following exposure of Apigenin to the Alzheimer’s neurons (+Aβ). Note that Apigenin and DA1 combination has a significant effect on enhancing expression of all three porosome proteins. **(Ba)** In this panel, the immunolocalization of SNAP-25 in untreated control neurons (Ctl), Aβ (Alzheimer’s) neurons, Aβ neurons treated with Apigenin (Aβ+Apigenin), Aβ neurons reconstituted with neuronal porosomes, Aβ treated with the DA1 peptide, Aβ treated with Apigenin and the DA1 peptide, and Aβ treated with porosome and the DA1 peptide, is shown. **(Bb)** In this panel, the immunolocalization of Syntaxin-1A in untreated control neurons (Ctl), Aβ (Alzheimer’s) neurons, Aβ neurons treated with Apigenin (Aβ+Apigenin), Aβ neurons reconstituted with neuronal porosomes, Aβ treated with the DA1 peptide, Aβ treated with Apigenin and the DA1 peptide, and Aβ treated with porosome and the DA1 peptide, is shown. **(Bc)** In this panel, the immunolocalization of AT1A3 in untreated control neurons (Ctl), Aβ (Alzheimer’s) neurons, Aβ neurons treated with Apigenin (Aβ+Apigenin), Aβ neurons reconstituted with neuronal porosomes, Aβ treated with the DA1 peptide, Aβ treated with Apigenin and the DA1 peptide, and Aβ treated with porosome and the DA1 peptide, is shown. **(Bd)** In this bar graph, the quantification of the immunolocalization of SNAP-25 in untreated control neurons (Ctl), Aβ (Alzheimer’s) neurons, Aβ neurons treated with Apigenin (Aβ+Apigenin), Aβ neurons reconstituted with neuronal porosomes, Aβ treated with the DA1 peptide, Aβ treated with Apigenin and the DA1 peptide, and Aβ treated with porosome and the DA1 peptide, is shown. **(Be)** In this bar graph, the quantification of the immunolocalization of Syntaxin-1A in untreated control neurons (Ctl), Aβ (Alzheimer’s) neurons, Aβ neurons treated with Apigenin (Abβ+Apigenin), Aβ neurons reconstituted with neuronal porosomes, Aβ treated with the DA1 peptide, Aβ treated with Apigenin and the DA1 peptide, and Aβ treated with porosome and the DA1 peptide, is shown. **(Bf)** In this bar graph, the quantification of the immunolocalization of AT1A3 in untreated control neurons (Ctl), Aβ (Alzheimer’s) neurons, Aβ neurons treated with Apigenin (Aβ+Apigenin), Aβ neurons reconstituted with neuronal porosomes, Aβ treated with the DA1 peptide, Aβ treated with Apigenin and the DA1 peptide, and Aβ treated with porosome and the DA1 peptide, is shown.

In early 2023, there were 187 Phase 1, 2, and 3 clinical trials assessing 141 drugs for AD. Thirty-six drugs were being assessed in Phase 3, 87 in Phase 2, and 31 in Phase 1. Although, neurotransmitter receptors, amyloid plaque, synaptic function, and inflammation were the most common targets of drugs in the pipeline, most therapies have focused on clearing the buildup of amyloid plaques and on reducing inflammation [36]. While these are important advancements in AD therapy, primarily focused on treatments to ameliorate the various consequences of the disease, there are no cures in sight nor are there therapies developed that treat the disease at the very early stage. In the current study, the systems level approach in correcting both the metabolic and secretory defects in AD at early stages of the disease, i.e., hyposmia or olfaction dysfunction during the preclinical stage of the disease [37], shows great promise. It is important to note that in >90% of Alzheimer’s patients, olfactory dysfunction precedes cognitive decline [38]. Moreover, the olfactory bulb of AD mice show loss in expression of SNAP-25 [39]. Dysfunction of odor discrimination can therefore be used as a predictive measure for AD [37], and the proposed [40] “Odorant Item Specific Olfactory Identification” test where certain odors can differentiate AD from general age- related decline in odor detection [40] should also be used in identifying patients for our ‘Metabolic and Secretory Corrector AD Therapy’. Both the DA1 peptide and Apigenin can pass through the blood-brain barrier. However, since the olfactory bulb is connected via nerves to the brain centers involved in learning, memory and emotion, the DA1 peptide combined either with Apigenin or the porosome, deliverable through the nasal route would be a viable and highly effective AD therapy, deliverable through the nasal route, especially at the early stages of the disease, would be a viable and highly effective AD therapy [Figure 4]. The therapy is scalable for commercialization since the DA1 circular peptide and Apigenin are readily available, and the scale-up of the neuronal porosome biologic could be achieved by immunoisolation from 3D cultures of the human neuroblastoma SH-SY5Y cell line grown in bioreactors [39]. 3D neuronal cultures will be used since the capacity of a neuron to establish synaptic connections is an essential prerequisite for its functional maturation, and that the formation of the synapse involves the appropriate assembly of the porosome secretory machinery at the nerve terminal [Figure 1A]. Since the neuronal porosome is present in all neurons, the above-mentioned scale-up approach offers reproducibility, reliability and safety. One needs to be critically aware that, since three [Figure 3] among possibly others, of the nearly 30 or so proteins of the neuronal porosome complex are impacted in Alzheimer’s, resulting in impairment function, one safe and highly effective approach would be to ameliorate the impairment by introducing normal neuronal porosomes at the pre-synaptic membrane. Porosome reconstitution therapy therefore remains the most optimal approach in ameliorating neurotransmitter release defects in AD. The normal secretion of neurotransmitter and its timely breakdown is physiologically relevant and optimal, as opposed to the available AD medications such as cholinesterase inhibitors that prevent the breakdown of acetylcholine in the brain to improve neuronal communication. Cholinesterase inhibitor drugs exhibit side effects that may include nausea, vomiting, diarrhea, muscle cramps, fatigue and even weight loss [40].

**Figure 4.**
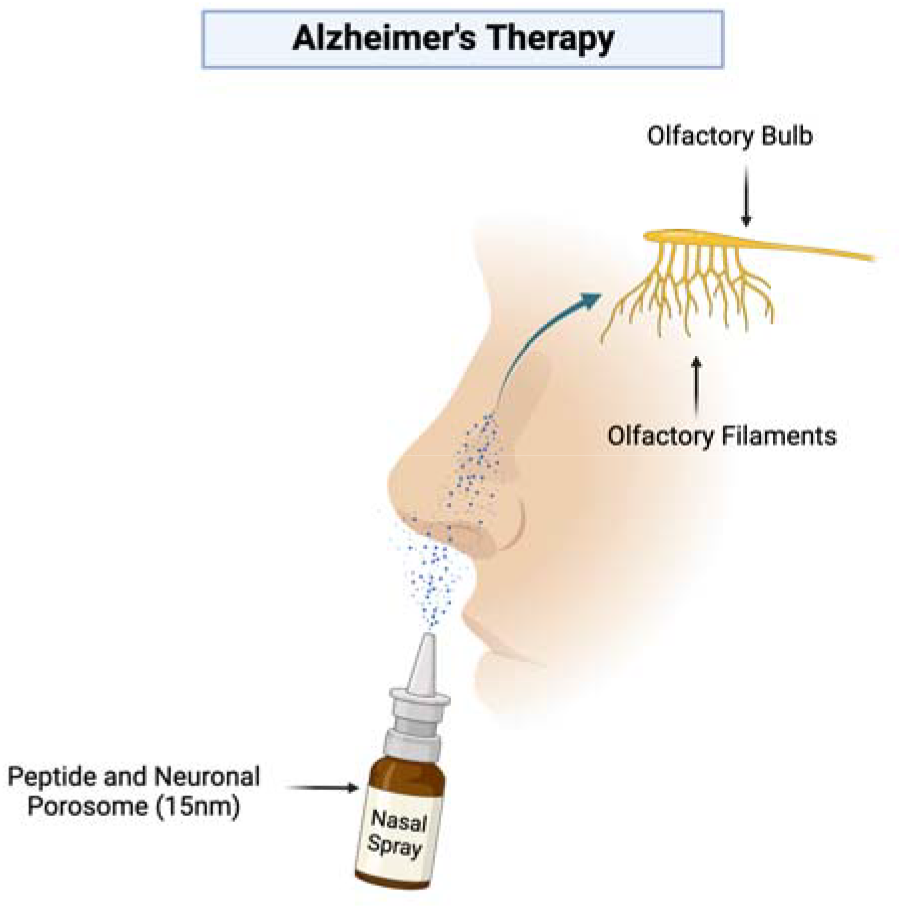
Combination therapy of neuronal porosome or the small flavonoid molecule Apigenin, in combination with the mitochondrial fission inhibiting DA1 peptide for AD therapy. Note that in >90% of Alzheimer’s patients olfactory dysfunction precedes cognitive decline and since the olfactory bulb is connected via nerves to brain regions involved in learning, memory and emotion, this suggests that a combinatorial DA1 peptide therapy either with Apigenin or porosome (a 15 nm biologic), delivered via the nasal route to reach the olfactory bulb, would be most effective in AD therapy, especially at the early stages of the disease.

## Contributions

The idea for the use of porosome-reconstitution therapy was developed at Porosome Therapeutics Inc. The research design was developed by B.P.J. and W-J.C, and conducted by YS under the expert guidance of XQ at Case Western Reserve University as paid contract from Porosome Therapeutics, Inc. B.P.J. wrote the paper. SP created the schematics, assembled the figures and formatted the manuscript. All authors participated in critical reading and discussion of the manuscript.

## Acknowledgements

Work presented in this article was supported by Porosome Therapeutics, Inc., and the Viron Molecular Medicine Institute, Boston, MA.

